# Impact of a tarsonemid prey mite and its fungal diet on the reproductive performance of a predatory mite

**DOI:** 10.1101/2020.10.14.338889

**Authors:** Dominiek Vangansbeke, Marcus V. A. Duarte, Jonas Merckx, Alfredo Benavente, Wojciech L. Magowski, Soraya C. França, Karel Bolckmans, Felix Wäckers

## Abstract

Phytoseiid predatory mites are the most important group of biocontrol agents currently implemented in protected cultivations worldwide. The possibility to produce these predators at high densities on factitious prey mites is a crucial factor for their success. Commonly used factitious prey mites comprise mainly species belonging to the cohort of Astigmatina. In the present study, we investigated the potential of tarsonemid prey mites as a food source for the spider mite predator *Neoseiulus californicus* (McGregor) (Acari: Phytoseiidae). The oviposition of *N. californicus* on mixed stages of *Tarsonemus fusarii* Cooreman (Acari: Tarsonemidae) was similar to that on its natural prey, the two-spotted spider mite *Tetranychus urticae* Koch (Acari: Tetranychidae). As most tarsonemids are specialized fungus-feeders, we tested the effect of different fungal species on the growth of *T. fusarii*. Subsequently, we analysed the impact on the fungal growing medium on the oviposition of *N. californicus*. The fungal growing medium of *T. fusarii* had a significant effect on the reproductive output of the predatory mite, having a negative effect. When *T. fusarii* was separated from the rearing medium, these detrimental effects were not observed. The present study shows the potential of using tarsonemid prey mites in the production of predatory mites.

## Introduction

Being one of the largest groups within the Heterostigmata, the family of Tarsonemidae currently consists of 43 genera and over 600 described species (Lofego et al., 2016). Mites from this family have a wide range of feeding behaviours, from true herbivores, algivores, fungivores to egg predators, and they occupy many natural (e.g. plants, soil litter) and human-made habitats (e.g. stored product facilities, plant cultivations) (Lindquist, 1986). Many species live in close association with insects, interacting either directly or indirectly with them. Direct interactions range from commensalism, phoresy to predation. Some species from the genus *Tarsonemus* compete with bark beetle larvae (e.g. *Dendroctonus* sp.) for fungal food, demonstrating indirect associations as well (Lombardero et al., 2003; Scott et al. 2008). For species, such as *Polyphagotarsonemus latus* (Banks), *Phytonemus pallidus* (Banks) and *Steneotarsonemus spinki* Smiley, which are destructive pests of a wide range of agricultural crops, numerous studies have investigated the biology of these species and their control (Easterbrook et al., 2001; Gerson, 1992; Hummel et al., 2009). In contrast, information is scarce on the biology and ecological role of fungivorous species, even though they comprise the majority of the tarsonemids (Lindquist, 1986). Species within the genus of *Daidalotarsonemus* and *Excelsotarsonemus* thrive on algae and fungi growing on the foliage of plants, predominantly in humid areas (Ochoa et al., 1995; Sousa et al., 2018). Fungi are also the primary food source of tarsonemids within the genus *Tarsonemus.* The cosmopolitan species *Tarsonemus fusarii* Cooreman was reported as a contaminant of fungal laboratory cultures and was shown to feed on a wide range of fungal species (Geeraerts, 1974; Gotz and Reichenberger, 1953). Fungal food preference was tested for *Tarsonemus confusus*, indicating that such preference exists, and specific fungus can alter the mites’ functional sex ratio (Belczewski and Harmsen 1997). Together with *Tarsonemus granarius* Lindquist, *T. fusarii* is also found in granaries thriving on a variety of fungi, such as *Chaetomium* and *Aspergillus* (Lindquist, 1972). *Tarsonemus fusarii* is one of the most ecologically widespread species (Kaliszewski & Sell 1990).

Phytoseiid predatory mites are important biocontrol agents used worldwide to control mite and insect pests (McMurtry et al., 2013). The possibility to mass-produce phytoseiids on factitious prey mites has been a major contributor to their successful implementation in biocontrol programs (Calvo et al., 2015; Ramakers and Van Lieburg, 1982). Traditionally, factitious prey mites belong to the cohort of Astigmatina (Zhang, 2003), such as *Carpoglyphus lactis* (L.) (Bolckmans and van Houten, 2006), *Lepidoglyphus destructor* (Schrank) (Castagnoli et al., 2006) and *Suidasia medanensis* (Oudemans) (Midthassel et al., 2013).

Phytoseiids effectively feed on both pest tarsonemids, such as *P. latus* (Duarte et al., 2015), and non-pest tarsonemids, such as *T. fusarii* (Vangansbeke et al., 2018). Here, we assessed the potential of *T. fusarii* to be used as a factitious prey mite in the production of the predatory mite *Neoseiulus californicus* (McGregor) (Acari: Phytoseiidae). The latter is typically used as a biological control agent of spider mites (McMurtry et al., 2013; van Lenteren, 2012).

For the production of astigmatid prey mites, mixes of bran, wheat germ, and yeast are commonly used food sources (Huang et al., 2013; Ramakers and Van Lieburg, 1982). As mouthparts of Tarsonemidae are stylet-like, adapted for piercing and sucking (Hughes, 1976; Kaliszewski et al., 1995), Nuzacci et al 2002), most species rely on the presence of fungal mycelia to enable food ingestion. Therefore, we explored two fungi that could be used as a food source for *T. fusarii*. Ideally, such fungal food sources should be 1) of high nutritional value for *T. fusarii*, 2) safe to use for biocontrol companies (absence of toxins) and 3) non-plant pathogenic. Two fungi, which should meet the latter two requirements, are *Aspergillus oryzae* and *Fusarium venenatum*. *A. oryzae* has been used for centuries in fermentation processes of rice and soy products (“koji”) in South-East Asia. It is generally accepted that *A. oryzae*, through years of cultivation, became a domesticated ecotype of *Aspergillus flavus* (Barbesgaard et al., 1992; Machida et al., 2005). During this domestication process, aflatoxins present in *A. flavus* have been eliminated (Domsch et al., 1980). *F. venenatum* is used for the production of mycoprotein for human consumption under the trade name “Quorn” (Wiebe, 2002). The strain A 3/5 (NRRL 26139 = ATCC 20334) used for commercial production does not produce aflatoxins (O’Donnell et al., 1998).

Here, we tested the nutritional value of *T. fusarii* for *N. californicus* as compared to its natural prey, the spider mite *Tetranychus urticae*. Secondly, we assessed different food sources for the growth of *T. fusarii*. For this, we compared a standard mix of wheat bran, wheat germ, and yeast with two fungi, *A. oryzae* and *F. venenatum*, grown on bran. Finally, the impact of *A. oryzae* on the reproductive performance of *N. californicus* was investigated.

## Material and Methods

### Mite colonies

Individuals of *Tarsonemus fusarii* were collected from *Rubus* sp. leaves in Gentbrugge (Belgium) (51.037472, 3.771090) and were maintained on a mixture of bran and wheat germ (50%/50%, w/w) at 22 ± 1 ^°^C, 80 ± 5 % relative humidity and a photoperiod of 16 hours light and 8 hours dark.

Two-spotted spider mites, *T. urticae*, were grown on bean plants in a separate greenhouse department at the greenhouse facilities of Biobest N.V. (Greenlab, Westerlo, Belgium).

*Neoseiulus californicus* females used for the experiments were collected from the mass-rearing facilities at Biobest N.V.

### Fungal strains

Both *Fusarium venenatum* Nirenberg (ATCC® 20334^™^) and *Aspergillus oryzae* var. *viridis* Murakami, anamorph (ATCC® 22788^™^) were grown on petri dishes containing potato dextrose agar (PDA Pronadisa). When the fungi were fully developed, circular pieces were removed from these agar Petri dishes to inoculate a mixture of equal parts of wheat bran and water with 5% (by weight) of potato dextrose broth (PDB Pronadisa). This mixture was incubated for 48 hours at 30°C at 90% R.H. before being offered to the mites. This mixture, as well as the combination of temperature, humidity, and time were previously shown to yield the best results for the mite rearings.

### Rearing of *T. fusarii* on bran inoculated with fungus

The mites were reared on each of the two fungal strains that had been grown on the wheat bran + PDB as described previously. The mites were kept at 22 ± 1 ^°^C, 80 ± 5 % relative humidity and a photoperiod of 16 hours light and 8 hours dark.

### Experiment 1: *T. fusarii* as a food source for *N. californicus*

In the first experiment, we performed a two-day egg-laying trial of *N. californicus* when provided with either *T. fusarii* or *T. urticae* as a food source. Young gravid female *N. californicus* were transferred individually from the mass-rearing medium on small PVC arenas (2 × 4 cm). Arenas were placed on wet cotton and the edges of the arena were covered with moist tissue paper to prevent the mites from escaping. Tarsonemid prey mites were offered *ad libitum* by transferring a small spoon of rearing medium (± 0.2 g) onto the PVC arena. For the spider mites, a small piece (1 × 1 cm) of heavily infested leaf was cut from the bean plants and was provided on the arena. Eggs produced by *N. californicus* were removed and counted daily. Per treatment, 12 replicates were set up. As the pre-experimental diet might affect the predator’s initial fecundity (Sabelis, 1990), the first day of egg-laying was omitted from the analysis. The average number of eggs produced on the second day were compared using GLM analysis with a Poisson distribution. Contrasts between diets were assessed by stepwise model simplification through aggregation of non-significant factor levels (Crawley, 2013). All statistical analyses were done using the computer software R version 3.6.1 (R Core Team 2014).

### Experiment 2: Growth of *T. fusarii* on different food sources

In this experiment, we compared the population growth of *T. fusarii* when fed on different food sources. These food sources were a bran inoculated with either a yeast (*Saccharomyces cerevisiae*) mix, or with one of the two fungi *A. oryzae* and *F. venenatum*. The food sources were prepared in the same manner as described above and were put in a 30ml insect breeding dish with a ventilated lid. Two thousand *T. fusarii* were added to two grams of each of the food sources and allowed to grow for ten days. At the end of these ten days, the number of mites in each of the food source was estimated by taking a sample (~ 0.100 g) and counting the total number of mobile mites and extrapolating to determine the total amount of mobile mites in each repetition. The average number of mobile *T. fusarii* were compared using GLM analysis with a normal distribution. Contrasts among treatments were determined with general linear hypothesis testing (function glht of the package lsmeans in R, Lenth 2016). All statistical analyses were done using the computer software R version 3.6.1 (R Core Team 2014).

### Experiment 3: Effect of *T. fusarii* rearing method on the oviposition of *N. californicus*

For this experiment, *T. fusarii* was reared on bran, which was previously inoculated with either *A. oryzae* or *F. venenatum* (see above). *T. fusarii* was grown on the fungal media for more than one month before the start of the experiment. Experimental procedures were similar, as was described for experiment 1. Only now, a small spoon (± 0.2 g) of *T. fusarii* grown on either bran with *A. oryzae* or *F. venenatum* was supplied on the arenas as a food source. Egg-laying of female predators was counted during four consecutive days. For each food source, 12 replicates were used. Again, the first day of oviposition was not used in the analysis. The difference between the food sources was compared using linear mixed-eJect models (LME) with diet, time, and their interaction as fixed factors. Individual mite was a random factor. Non-significant interaction and factors were removed until a minimal satisfying model was reached (Crawley, 2013).

### Experiment 4: Effect of *T. fusarii* rearing medium on the oviposition of *N. californicus*

Here, we tested whether the *A. oryzae* rearing medium of *T. fusarii* could affect the reproductive performance of *N. californicus*. Therefore, we provided the predatory mite females with *T. fusarii* that were either sieved out from the fungus/bran medium or provided together with the fungus/bran mix. *T. fusarii* were separated from the rearing medium using a 100µm sieve. Hence, besides *T. fusarii*, also ample spores and short pieces of mycelia were present in the sieved out material. The trial set-up and statistical analysis was similar as was described for experiment 2.

## Results

### Experiment 1: *T. fusarii* as a food source for *N. californicus*

Diet did not affect the oviposition of *N. californicus* (GLM: Chi^²^=17.939; df= 1, 18; p=0.729), with 1.8 ± 0.3 eggs/female/day and 1.6 ± 0.4 eggs/female per day when offered *T. urticae* and *T. fusarii*, respectively.

### Experiment 2: Growth of *T. fusarii* on different food sources

There was a significant effect of the different food sources on the total number of mites (GLM: F=7.1; df= 2, 16; p>0.01). The number of *T. fusarii* was greater when reared on *A. oryzae* as compared to the other food sources (Figure 1). There were no differences between the total number of mites between the mix with yeast and *F. venenatum* treatments.

**Fig1:**
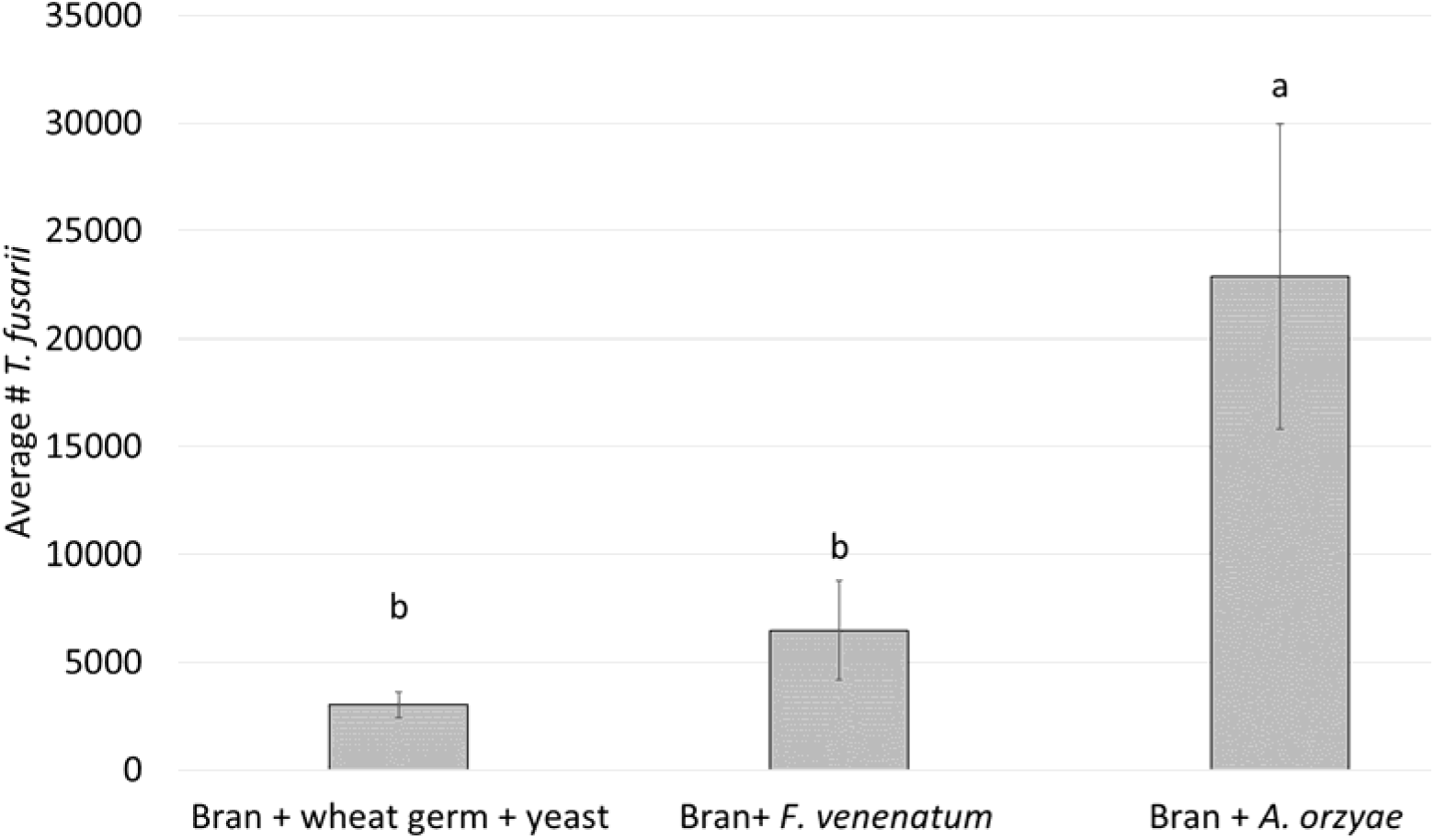
Mean (± standard error) number of mobile *T. fusarii* grown for 10 days on a mix of bran, wheat germ and yeast, or bran, PDB and *F. venenatum*, or *A. oryzae*. Initial population: 2000 mobiles. Bars with different letters differ significantly (contrasts after GLM, p < 0.05)

### Experiment 3: Effect of *T. fusarii* rearing method on the oviposition of *N. californicus*

There was no significant effect of the interaction between time and diet (LME: Chi^²^=3.394; df= 1; p=0.494) and time (LME: Chi^²^=1.455; df= 1; p=0.483). Diet had a significant effect on the egg-production of *N. californicus* (Figure 2, LME: Chi^²^=23.234; df= 1; p<0.0001), with a twofold increase in oviposition when fed *T. fusarii* that was produced on *F. venenatum* as compared to mites produced on *A. oryzae*.

**Fig2:**
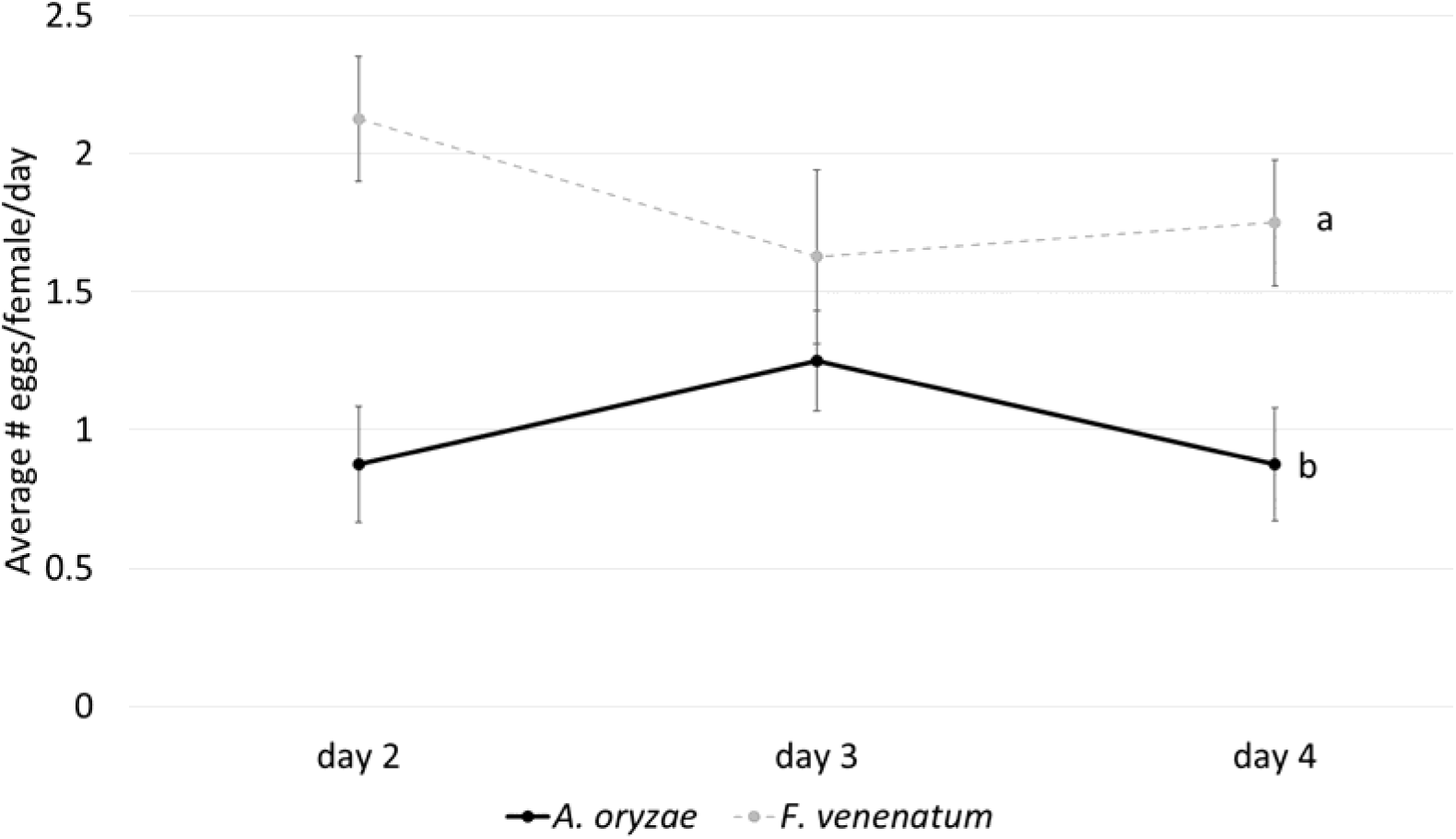
Mean (± standard error) number of eggs produced by *N. californicus* when fed *T. fusarii* either reared on bran with *A. oryzae* or *F. venenatum* (contrasts after LME, p < 0.05)

**Fig3:**
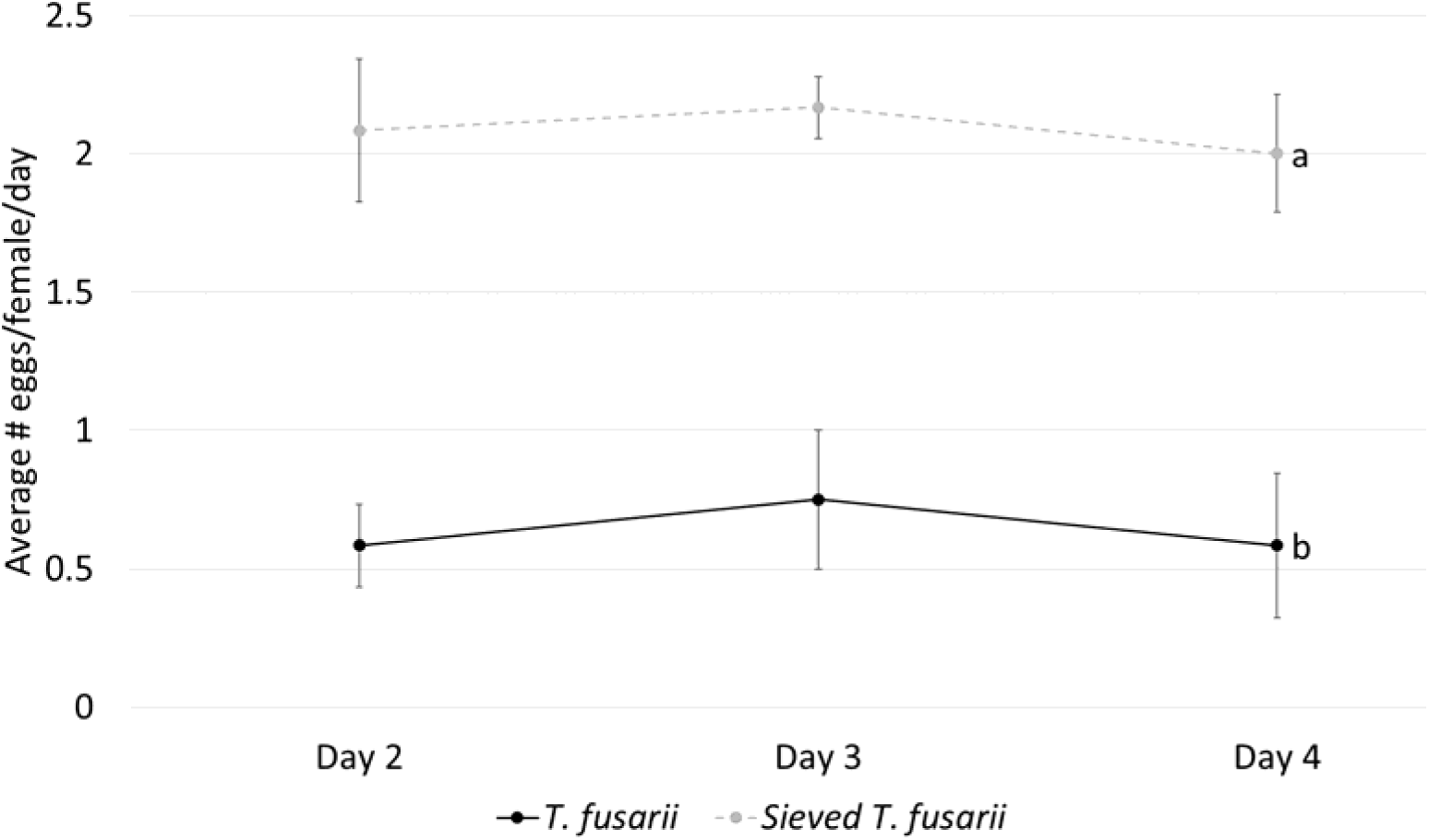
Mean (± standard error) number of eggs produced by *N. californicus* when fed *T. fusarii* either provided with the *A. oryzae*/bran medium or sieved out from it (contrasts after LME, p < 0.05)

### Experiment 4: Effect of *T. fusarii* rearing medium on the oviposition of *N. californicus*

Time and the interaction between diet and time did not have a significant effect (time= LME: Chi^²^=0.705; df= 1; p=0.703 and diet x time= LME: Chi^²^=0.054; df= 1; p=0.973). Sieving out *T. fusarii* significantly increased the oviposition of *N. californicus* (**Error! Reference source not found.**, LME: Chi²=37.069; df= 1; p<0.0001).

## Discussion

Augmentative biological control is a key component in integrated pest management, especially in protected cultivation of vegetable and ornamental crops (Naranjo et al., 2015). Phytoseiid predatory mites comprise the most important group of commercially available natural enemies (van Lenteren, 2012). An important factor explaining this success is the possibility to mass-produce these predatory mites at high densities on factitious astigmatid prey (Calvo et al., 2015; Knapp et al., 2018). However, further improvement of the efficiency of mass-production systems remains essential to reliably supply growers with an economic product (Knapp et al., 2018) and to further expand the scope towards open-field applications of biocontrol agents.

Many phytoseiids are known to feed, reproduce and control pest tarsonemids, such as the broad mite *P. latus* (Badii and McMurtry, 1984; Duarte et al., 2015; Fan and Petitt, 1994), the cyclamen mite *P. pallidus* (Easterbrook et al., 2001; Tuovinen and Lindqvist, 2010) and *Steneotarsonemus* spp. (Messelink and van Holstein-Saj, 2007; Zhang, 2003). Studies on feeding behaviour of phytoseiids on non-pest tarsonemids, however, are scarce. Only recently, *Neoseiulus barkeri* Hughes was reported to feed on *Tarsonemus confusus* Ewing (Li et al., 2018). For *Amblyseius swirskii* Athias-Henriot, feeding on *T. fusarii* resulted in an equally high oviposition rate as compared to feeding on the astigmatid *C. lactis* (Vangansbeke et al., 2018). In this study, also *T. fusarii* proved to be a good food source for the spider mite predator *N. californicus*.

Mites from the genus *Tarsonemus* are considered generalist fungus-feeders (Lindquist, 1986). Some species, such as *Tarsonemus ips* Lindquist and *Tarsonemus krantzi* Smiley & Moser, use their C-tergite ventral flaps to carry fungal spores (Moser, 1985). Other species are associated with feeding on fungal diseases, such as *T. confusus* feeding on *Cladosporium* sp. and *Alternaria* sp. (Belczewski and Harmsen 1997; Van der Walt et al., 2011). Although some instances have been reported where *Tarsonemus* representatives were observed to be associated with plant damage in some ornamental crops, it is believed that this species, and in extension all mites within the genus of *Tarsonemus*, are fungivorous and are never the causal agents of primary plant damage (Zhang, 2003).

Occasionally, *Tarsonemus* species have been reported as pests in mushroom cultivation (*Tarsonemus myceliophagus* (Hussey and Gurney, 1967)) or stored grain facilities (*Tarsonemus granarius* (Lindquist, 1972)).

Here, *A. oryzae* grown on bran was a superior food for *T. fusarii*, with seven-fold larger population numbers as compared to that on a standard astigmatid growing medium consisting of wheat bran, wheat germ and yeast. Although not significantly different, *F. venenatum* also resulted in higher numbers of tarsonemid prey mites. Several factors determine the acceptance and subsequent performance of a mite feeding on fungus. Besides the ability to digest the ingested fluids (Hubert et al., 2001), the presence of mycotoxins potentially affects the suitability of fungal food source (Parkinson et al., 1991; Rodriguez et al., 1980). As both fungi tested are considered mycotoxin-free, the latter reason seems unlikely. Another explanation for the superiority of *A. oryzae* over *F. venenatum* might be the difference in the production of fungal spores. The *Aspergillus* strain used in our experiments produced an abundance of spores as opposed to the *F. venenatum* fungus, which was not sporulating under the test conditions. The stylet-like mouthparts of *T. fusarii* are equipped with apodemes (being attachment structures for muscles), which are one of the taxonomic determinants for this species. Hence, *T. fusarii* might be adapted to feed on fungal spores as well as fungal mycelium (R. Ochoa, personal communication), which could explain the strong growth response to *A. oryzae*.

While both *A. oryzae* and *F. venenatum* were shown to be suitable food source for growing tarsonemids, other fungi that are preferably mycotoxin-free, could also be considered when producing prostigmatid and astigmatid prey mites used for the production of natural enemies. Several fungal genera, such as *Chaetomium*, *Aspergillus, Trichoderma* and *Alternaria* are associated with tarsonemids (Smiley and Moser, 1985; White and Sinha, 1981). Alternatively, other food-safe fungi, like *Rhizopus sp*., *Actinomucor* sp., *Amylomyces* sp., *Mucor* sp., *Monascus* sp., could be tested for their suitability to rear prey mites. However, *Penicillium* sp, was proven to generate detrimental effect onto the developement of some *Tarsonemus* species populations (Suski, 1972).

It is unclear why the oviposition of *N. californicus* decreased in the presence of the *A. oryzae* growing on bran. Some hypotheses explaining this phenomenon are:

1. *A. oryzae* growing on bran produces volatiles that negatively impact the predator. When growing on rice, *A. oryzae* has been shown to produce a range of volatile compounds, including aldehydes and ketones. Moreover, production of alcohols and aldehydes increased under more anaerobic conditions (Ito et al., 1990). As *N. californicus* was frequently found hiding inside pieces of bran, oxygen deficiency might have caused a more toxic microenvironment resulting in the observed negative outcome.
2. When a fungus such as *A. oryzae* is growing on bran, thereby producing an abundance of spores, prey mites are more difficult to find as compared to sieved our material. Habitat complexity has previously shown to affect both the movement of the predator and their prey capture rate (Lee and Zhang, 2016; Madadi et al., 2007). Hence, when reared on *A. oryzae* on bran, the habitat structure might have impeded *N. californicus* in successfully locating and subduing the tarsonemid prey.
3. Fungus growing on the bran substrate might have altered the microclimate, which negatively affects the phytoseiid’s performance. Growth of micro-organisms, including fungi, is associated with the production of heat (van Kleeff et al., 1993; Wilson, 1999). As predatory mites are ectothermic organisms with a maximum temperature threshold (Vangansbeke et al., 2015), excessive heat production can result in deleterious effects.

In summary, we demonstrate the potential of using tarsonemid prey mites in the production of the phytoseiid predatory mite *N. californicus*. The impact of the fungal species and its growing media on both the tarsonemid prey mite and *N. californicus* merits further investigation.

## Acknowledgements

We would like to thank Peggy Bogaerts and Ilse Jacobs for the help with growing the plant material and maintenance of the spider mite production.

